# N-(3-oxododecanoyl)-L-homoserine lactone interactions in the breast tumor microenvironment: implications for breast cancer viability and proliferation *in vitro*

**DOI:** 10.1101/132092

**Authors:** Brittany N. Balhouse, Logan Patterson, Eva M. Schmelz, Daniel J. Slade, Scott S. Verbridge

**Affiliations:** School of Biomedical Engineering and Sciences, Virginia Tech-Wake Forest University, Blacksburg, VA, United States of America; Department of Biomedical Engineering and Mechanics, Virginia Tech, Blacksburg, VA, United States of America; Department of Biochemistry, Virginia Tech, Blacksburg, VA, United States of America; Department of Pathology, University of Virginia, Charlottesville, VA, United States of America; Department of Human Nutrition, Foods and Exercise, Virginia Tech, Blacksburg, VA, United States of America

## Abstract

It is well documented that the tumor microenvironment profoundly impacts the etiology and progression of breast cancer, yet the contribution of the resident microbiome within breast tissue remains poorly understood. Tumor microenvironmental conditions, such as hypoxia and dense tumor stroma, predispose progressive phenotypes and therapy resistance, however the role of bacteria in this interplay remains uncharacterized. We hypothesized that the effect of individual bacterial secreted molecules on breast cancer viability and proliferation would be modulated by these tumor-relevant stressors differentially for cells at varying stages of progression. To test this, we incubated human breast adenocarcinoma cells (MDA-MB-231, MCF-DCIS.com) and non-malignant breast epithelial cells (MCF-10A) with N-(3-oxododecanoyl)-L-homoserine lactone (OdDHL), a quorum-sensing molecule from *Pseudomonas aeruginosa* that regulates bacterial stress responses. This molecule was selected because *Pseudomonas* was recently characterized as a significant fraction of the breast tissue microbiome and OdDHL is documented to impact mammalian cell viability. After OdDHL treatment, we demonstrated the greatest decrease in viability with the more malignant MDAMB-231 cells and an intermediate MCF-DCIS.com (ductal carcinoma *in situ*) response. The responses were also culture condition (i.e. microenvironment) dependent. These results contrast the MCF-10A response, which demonstrated no change in viability in any culture condition. We further determined that the observed trends in breast cancer viability were due to modulation of proliferation for both cell types, as well as the induction of necrosis for MDA-MB-231 cells in all conditions. Our results provide evidence that bacterial quorum-sensing molecules interact with the host tissue environment to modulate breast cancer viability and proliferation, and that the effect of OdDHL is dependent on both cell type as well as microenvironment. Understanding the interactions between bacterial signaling molecules and the host tissue environment will allow for future studies that determine the contribution of bacteria to the onset, progression, and therapy response of breast cancer.

## Introduction

The tumor microenvironment is now a widely recognized and well-studied contributor to cancer dynamics, particularly for breast cancer. While increased matrix density, programming of cancer-associated stromal cells, evolving gradients of oxygen and nutrients, and leaky vasculature have all been implicated as key players in breast cancer progression (1-4), the impact of the recently identified breast tissue resident microbiotic niche has received little attention. Beyond the effects of pathogenic or tumorigenic bacteria such as *Chlamydophila pneumonia*, *Salmonella typhi*, *Streptococcus gallolyticus* (5), *Helicobacter pylori (6)* and *Fusobacterium nucleatum* (7), the majority of analyses of tumor-microbiome interactions have centered on local cell-cell interactions within the gut microenvironment, or more systemic immune effects influenced by gut microbiota (8). Only a handful of studies have been conducted to investigate the influences of tissue-resident bacteria in other tumor sites, such as for breast cancer (9-11). Even fewer have investigated how small molecules released from resident bacteria may interact with cells in the presence of other critical microenvironmental factors, e.g. tumor hypoxia, to regulate cancer progression. In an effort to address these questions, we investigated interactions between the quorum-sensing molecule N-(3-oxododecanoyl)-L-homoserine lactone (OdDHL) and the breast tumor relevant microenvironmental cues of a stiff collagen-derived tissue mimic and hypoxia. This representative study will aid in our understanding of how the understudied breast tissue microbiome may contribute to disease phenotypes, patient-to-patient variability, and cancer progression.

OdDHL secreting *Pseudomonas* are Gram-negative Proteobacteria that were recently found to make up a significant fraction of the microbiome within breast tissue (12), and on nipple skin and aspirate from women with and without a history of breast cancer (13). In addition, *Pseudomonas* has been shown to make up a significant portion of the bacteria found in human breast milk (11, 14-16). *Pseudomonas* was among the top five most abundant genera for both sample populations in the 2014 study by Urbaniak et al. (12) and in several studies of breast milk microbiota (14). OdDHL is a quorum-sensing molecule associated with biofilm development and environmental stress response in bacteria (17) that has been shown to promote apoptosis in a variety of human cell lines (18-23). Especially interesting is its selective effect inhibiting proliferation and inducing apoptosis in breast cancer cells, but not in non-malignant breast cells (21). For these reasons, studying the effects of OdDHL is not only important from the basic science perspective (e.g. regarding its role in the tumor microenvironment), but also from the therapeutic perspective given its potential as an anti-cancer treatment (24, 25).

High tissue density is not only a risk factor for breast cancer development, but also a clinical diagnostic tool (26). Increased stroma density (resulting from increased collagen deposition and cross-linking in the extracellular matrix (ECM)) is also associated with phenotypic changes in both cancer cells and normal stroma cells (27). In an *in vitro* setting, growing cells on or within a collagen hydrogel, as opposed to on a polystyrene substrate in two-dimensions (2D), changes not only the type of focal adhesions cells make with their environment (28), but also how cells respond to stresses within that environment (e.g. chemotherapeutic agents) (27, 29, 30). Three-dimensional (3D) cellular adhesions are implicated in reduced cellular response to chemotherapeutics (a phenomenon called adhesion-mediated resistance) and a progressive phenotype (27, 31, 32). Hypoxia is also a well-documented constituent of the solid tumor microenvironment; angiogenesis cannot keep up with the rate of tumor growth and produces a gradient of oxygen from the well-fed periphery to a necrotic core that is devoid of oxygen (33). Similar to cancer cells in a 3D environment, a low oxygen environment is associated with a progressive phenotype and a resistance to chemotherapy (33, 34). Although, adhesion- and hypoxia-mediated resistance have been extensively demonstrated for chemotherapeutic compounds, adhesion- and hypoxia-mediated regulation of cellular response to bacterial signaling molecules has not yet been investigated. Despite the previous research into OdDHL, the role of OdDHL in the context of breast tumor microenvironmental stressors was still unknown. Because such stressors are associated with altered cell responses (e.g. chemoresistance), it was our hypothesis that OdDHL would differentially modulate breast cancer cell viability and proliferation in hypoxia and in 3D culture (in which cells are seeded in the bulk of a tissue mimic).

In an effort to make a preliminary characterization of the response for both metastatic and non-metastatic subtypes of breast cancer, we utilized MDA-MB-231 and MCF-DCIS.com cells, respectively. MCF-10A breast epithelial cells were used as a non-malignant control. Both MDAMB-231 and MCF-10A cells were chosen for this research because of their wide documentation in the literature (35, 36). The MCF-DCIS.com cell line, which is derived from the MCF-10A line (37, 38), offers an intermediary model cell between the MCF-10A and MDA-MB-231 lines; not only does it represent a pre-invasive, non-metastatic form of breast cancer, but is Her2+ (37) whereas the MDA-MB-231 line is triple negative (35). In addition, both necrotic cores and desmoplasia have been observed in lesions from MCF-DCIS.com injections (38) as well as in MDA-MB-231 xenografts (39, 40), motivating our studies in hypoxia- and adhesion-mediated resistance.

## Methods

### Cell culture

We used MDA-MB-231 triple negative breast cancer cells (ATCC® HTB-26™), MCF-DCIS.com ductal carcinoma *in situ* breast cancer cells (generously donated by Dr. Eva M. Schmelz) (38), and MCF-10A breast epithelial cells (ATCC® CRL-10317™) for our malignant versus non-malignant cell comparison. MDA-MB-231 cells were cultured in DMEM/F12 (1:1) media supplemented with 10% fetal bovine serum and 1% penicillin-streptomycin (PS). MCF-10A and MCF-DCIS.com cells were cultured in DMEM/F12 (1:1) media supplemented with 5% horse serum, 1% PS, 0.05% hydrocortisone, 0.1% human insulin, 0.02% epidermal growth factor, and 0.01% cholera toxin. Hereafter, these supplemented DMEM/F12 (1:1) media will be referred to as complete media. Cells were subcultured every 4-5 days and used until passage 50. Stock cells were grown in T-75 flasks under standard culture conditions (37°C incubator with 5% CO_2_, and ambient O_2_ (referred to hereafter as normoxia).

For 2D experimental conditions, cells were seeded in 48-well polystyrene plates at a concentration of 100,000 cells per milliliter (30,000 cells per well), with two plates seeded per cell type per experiment. After seeding, all plates were incubated under standard culture conditions in normoxia. After 24 hours, the media was changed in all wells and the plates that were to undergo the hypoxia treatment were moved into an incubator with 1% oxygen and the standard culture conditions of 37°C, 5% CO_2_.

### 3D culture

3D culture was defined as cells suspended throughout a tissue mimic, namely a rat tail derived type I collagen hydrogel. The collagen stock was manufactured using a technique described previously (41). Briefly, collagen was extracted from Sprague-Dawley rat tails (Bioreclamation, Inc), sterilized in 70% ethanol, and dissolved in 0.1% glacial acetic acid for three days. The dissolved collagen stock was then aliquoted, frozen, lyophilized, and stored at −20°C. At least three days prior to use, the lyophilized collagen was reconstituted in 0.1% glacial acetic acid at a concentration of 12 milligrams of collagen per milliliter of acetic acid solution, and this dissolved stock solution was then used for up to four weeks post re-constitution.

The hydrogel was created by neutralizing the collagen stock with empirically derived proportions of 10X DMEM, 1X DMEM/F12 (1:1) and 1M sodium hydroxide (NaOH) (revised from (41)) to create a hydrogel with pH 7.4 and 6 mg/mL of collagen in 0.1% acetic acid solution. During the neutralization process, all solutions were kept on ice to prevent collagen polymerization. Cells were suspended throughout the hydrogel by resuspension of a cell pellet into the required 1X DMEM/F12 (1:1) prior to its addition into the unpolymerized (but neutralized) hydrogel solution. Cells were seeded in the hydrogel solution at a concentration of three million cells per milliliter of solution (~300,000 cells per well). The hydrogel solution was pipetted into pretreated SYLGARD® 184 silicone elastomer molds topped with untreated coverslips to create approximately 1 mm thick discs comprised of ~0.1 mL hydrogel solution. The hydrogels were incubated for 30 minutes at 37°C, then transferred into 24-well plates and covered with 0.6 mL complete media standard of the included cell type. There was one plate of 3D samples per cell type. All 3D samples were maintained in an incubator under standard culture conditions (37°C, 5% CO_2_, normoxia). Hydrogels with no cells were also created as a control for the alamarBlue™ cell viability assay described below.

### OdDHL treatment

Two days after seeding, the appropriate complete media containing varying concentrations of OdDHL were added to the samples. The concentrations of OdDHL reflect levels documented in the literature as physiological values in a niche in close proximity to the bacteria producing the quorum sensing molecule (42). For the OdDHL stock solution, lyophilized OdDHL (Sigma-Aldrich) was reconstituted in dimethyl sulfoxide (DMSO) at a concentration of 67.25 mM or 20 mg/mL (the maximum solubility of OdDHL in DMSO). For the experimental solutions, the OdDHL stock solution was added to the complete medium appropriate for the cell line (as described above) to create the highest concentration (400 μM) OdDHL solution. Serial 1:2 dilutions of the stock solution were performed in DMSO to create 200 μM, 100 μM, 50 μM, and 25 μM solutions in complete medium. A control of 0.6% DMSO (the volume per volume percentage of DMSO in all OdDHL treatments) was used. Cells were treated with the OdDHL or control solutions for 24 hours before proliferation (by EdU assay) and apoptosis/necrosis (by annexin V and propidium iodide assay) was analyzed, and 48 hours before viability (by alamarBlue™ assay) was analyzed. The difference in incubation period was determined by the nature of the assay to be performed.

### alamarBlue™ cell health assay

After the OdDHL or control treatment, the treatment solutions were removed and fresh complete medium supplemented with 10% alamarBlue™ solution (Thermo Fisher Scientific) was added to all wells, including 2D and 3D negative control wells where there were no cells. These were incubated in normoxic or hypoxic incubators until positive control wells (0.6% DMSO control) were magenta in color; reduction rate was dependent on cell type and culture condition. When controls wells showed the color change, three subsamples were taken from the solution of each well and added to a 96-well plate. As per the manufacturer’s instructions, the 96-well plates were read on a spectrophotometer, which analyzed the absorbance at 570 and 600 nm wavelengths. The percent reduction of alamarBlue™ was calculated with the equation recommended by the manufacturer, namely:

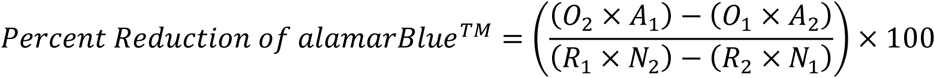

where *O*_1_ is the molar extinction coefficient (E) of oxidized alamarBlue™ at 570 nm, *O*_2_ is the E of oxidized alamarBlue™ at 600 nm, *R*_1_ is the E of reduced alamarBlue™ at 570 nm, *R*_2_ is the E of reduced alamarBlue™ at 570 nm, *A*_1_ is the absorbance of the test well at 570 nm, *A*_2_ is the absorbance of the test well at 600 nm, *N*_1_ is the absorbance of the negative control at 570 nm and *N*_2_ is the absorbance of the negative control at 600 nm. The average percent reduction of alamarBlue™ for each experimental group was divided by the average percent reduction of the 0 μM OdDHL (0.6% DMSO) control to calculate a relative viability.

### EdU assay

An EdU assay was performed to study the effect of OdDHL on proliferation of cells in normoxia (2D), hypoxia (2D), and 3D (normoxia). The EdU assay was conducted using the Click-iT® EdU Alexa Fluor 488® Imaging Kit (Thermo Fisher Scientific) as per the manufacturer’s instructions, with the exception of the fixing where 10% formalin was substituted for 3.7% formaldehyde. Briefly, after incubation with OdDHL in the various experimental conditions for 24 hours, half of the experimental solution was removed and fresh complete medium supplemented with the EdU molecule was added to each well (including controls). After a two hour incubation, the samples were fixed with 10% formalin (15 minutes for 2D, 45 minutes for 3D), washed and incubated with the Click-iT® reaction cocktail to fluorescently label cells that entered the S-phase of the cell cycle during the two hour incubation period. Cells were then counterstained with DAPI. Wells were imaged under 100X magnification with 1 subsample per well for 2D conditions and, using a confocal microscope, 5 subsamples per well for 3D conditions vertically through the hydrogel volume (30 μm spacing). The images were post-processed in ImageJ; see Supporting Information for more information on post-processing (Fig S1). The area of cell nuclei stained with Alexa Fluor 488® was divided by the area of cell nuclei stained with DAPI to find the fraction of cells entering S-phase during the 2 hour incubation period.

### Apoptosis/necrosis assay

The apoptosis and necrosis response was determined by incubation with annexin V-FITC (AV-FITC) and propidium iodide (PI) using a commercial assay (Annexin V-FITC Apoptosis Kit, BioVision Inc.). The assay was completed according to manufacturer instructions for all 2D samples and with three times the recommended incubation time for 3D samples. After incubation, cells were immediately imaged using 100X magnification with 3 subsamples per well for 2D conditions and, using a confocal microscope, 5 subsamples per well for 3D conditions vertically through the hydrogel volume (30 μm spacing). The images were post-processed in ImageJ; see Supporting Information for more information on post-processing (Fig S2). All values are relative to the appropriate 0.6% DMSO control for each experimental condition.

### Statistical analysis

For each experiment, there were two wells per experimental condition and control and each experiment was conducted three times (N = 3), unless otherwise noted. The statistics are based on the average values for each condition in each experiment.

The statistical analyses consisted of a full factorial two-way analysis of variance (ANOVA) in which the differences in viability (relative percent reduction of alamarBlue™), proliferation rate (percent of cells entering S-phase), and apoptosis/necrosis (AV-FITC/PI stained cells) for varying culture conditions and OdDHL concentrations were analyzed for each cell line tested. Where a p-value < 0.05 was found for the ANOVA, post-hoc Tukey tests and least means contrasts were performed to analyze the degree of significance between experimental conditions. Error bars represent standard error of the mean.

## Results

We first compared highly malignant MDA-MB-231 cells with non-malignant MCF-10A cells. We found that the response to OdDHL was dependent not only on cell-type, but also the culture condition. As compared to the control, the malignant MDA-MB-231 cells showed significantly different responses to 400 μM of OdDHL in all culture conditions (Fig 1A-1E). In the 2D/normoxia condition, the 400 μM OdDHL treatment corresponded to approximately 52.2% (± 2.2%) viability, relative to the control. In the 2D/hypoxia condition, that relative viability was increased to 60.6% (± 2.2%) and in 3D/normoxia condition, that viability was increased to 81.9% (± 2.2%) (Fig 1C and Fig 1D, respectively). There were significant decreases in the MDA-MB-231 viability at the 100 and 200 μM levels of OdDHL, as compared to the control, for the 2D conditions as well (Fig 1A and Fig 1B). In contrast, the non-malignant MCF-10A cells exposed to OdDHL showed no significant change in viability relative to the control across all concentrations in all culture conditions (Fig 1F-1H).

**Figure 1.**
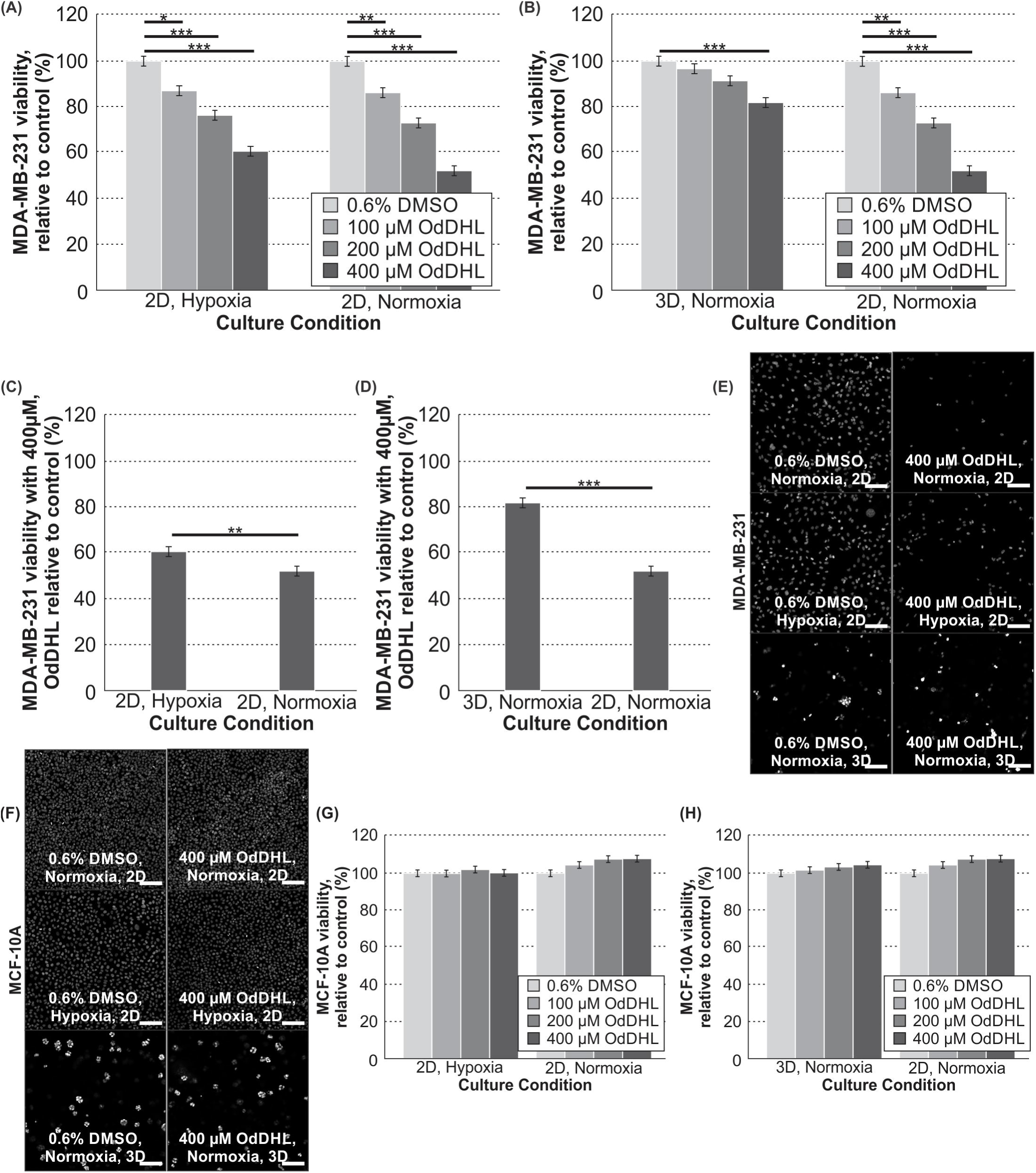
OdDHL has a cell-specific and significant impact on MDA-MB-231 viability in all culture conditions. Significant differences in mean MDA-MB-231 viability at 100, 200 and 400 μM concentrations relative to the control in hypoxia and normoxia in 2D (A) and 3D and 2D in normoxia (B); *** = p-value < 0.0001, ** = p-value < 0.01, * = p-value < 0.05 based on Tukey post-hoc differences. Significant differences in MDA-MB-231 viability relative to the control between culture conditions with 400 μM treatment in hypoxia and normoxia in 2D (C) and 3D and 2D in normoxia (D); *** = p-value < 0.0001, ** = p-value < 0.01 based on least mean contrasts. (E) Photomicrographs of MDA-MB-231 cells cultured in normoxia (2D), hypoxia (2D), and 3D (normoxia) with fluorescence microscopy (with DAPI staining; scale bar = 100 μm) at 100X magnification after 24 hours of OdDHL treatment. (F) Photomicrographs of MCF-10A cells cultured in normoxia (2D), hypoxia (2D), and 3D (normoxia) with fluorescence microscopy (with DAPI staining; scale bar = 100 μm) at 100X magnification after 24 hours of OdDHL treatment. Differences in MCF-10A viability relative to the control at 100, 200 and 400 μM concentrations in hypoxia and normoxia in 2D (G) and 3D and 2D in normoxia (H). Error bars represent standard error of the mean.

We next sought to quantify the viability response of the MCF-DCIS.com cells, which represent an intermediate phenotype between the non-malignant and metastatic extremes. The viability assay of the MCF-DCIS.com cells showed the cells responded in a similar fashion to MDA-MB-231 cells in both the 2D/normoxia and 2D/hypoxia conditions, with significant dose-dependent decreases in viability with increasing OdDHL concentrations (Fig 2A). However, unlike the MDA-MB-231 cells, the MCF-DCIS.com cells did not have a significant decrease in viability in the 3D/normoxia condition at any treatment level of OdDHL (Fig 2B); this is an interesting observation to which we will return shortly. Also, unique to the MCF-DCIS.com cells is a significantly lower viability at the 400 μM level for MCF-DCIS.com cells cultured in 2D/hypoxia than those cultured in 2D/normoxia (Fig 2C). However, for MCF-DCIS.com cells grown in 3D versus 2D in normoxia, there was significantly increased MCF-DCIS.com viability with the 400 μM OdDHL treatment (Fig 2D). An analysis of the cell specific viability response to 400 μM OdDHL showed that MCF-DCIS.com cells have significantly higher viability than MDA-MB-231 cells, but significantly lower viability than MCF-10A cells when cultured under the 2D/normoxia condition (Fig 2E). With the same treatment, MCF-DCIS.com cells showed significantly lower viability than MCF-10A cells when cultured under the 2D/hypoxia (a response that is statistically not different from that of MDA-MB-231 cells under the same condition) (Fig 2E). When cells were cultured under the 3D/normoxia condition, there was no difference in MCF-DCIS.com viability as compared to MCF-10A cell viability with the 400 μM OdDHL treatment; both were significantly higher than MDA-MB-231 cell viability under the same condition (Fig 2F).

**Figure 2.**
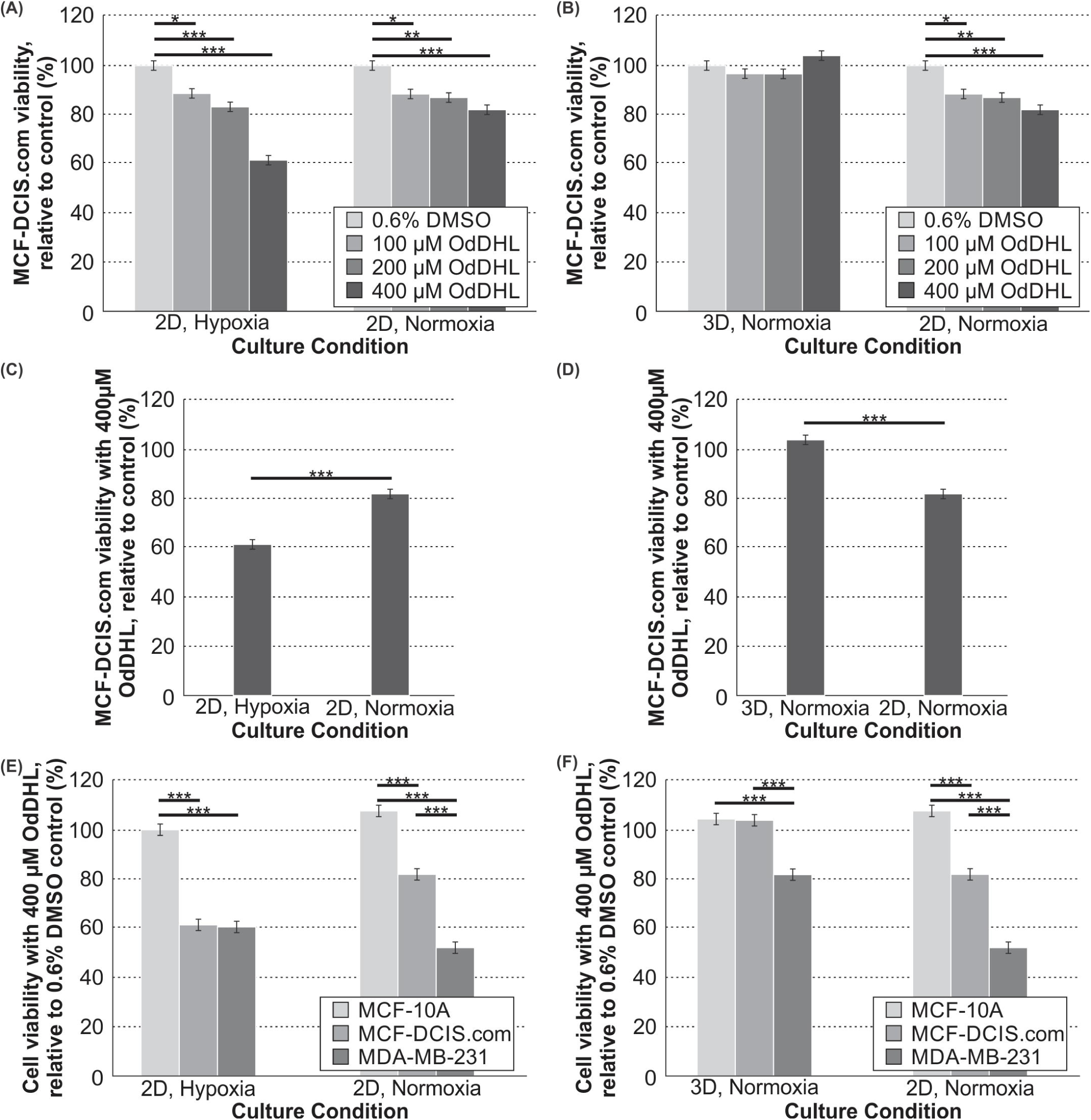
OdDHL has a cell-specific and differential impact on MCF-DCIS.com cells dependent on culture condition. Significant differences in mean MCF-DCIS.com viability relative to the control at 100, 200 and 400 μM OdDHL concentrations in hypoxia and normoxia in 2D (A) and 3D and 2D in normoxia (B); *** = p-value < 0.0001, ** = p-value < 0.01, * = p-value < 0.05 based on Tukey post-hoc differences. Significant differences in MCF-DCIS.com viability relative to the control between culture conditions with 400 μM OdDHL treatment in hypoxia and normoxia in 2D (C) and 3D and 2D in normoxia (D); *** = p-value < 0.0001 based on least mean contrasts. Significant differences in mean MCF-10A, MCF-DCIS.com, MDA-MB-231 viabilities relative to the control at 400 μM OdDHL concentrations in hypoxia and normoxia in 2D (E) and 3D and 2D in normoxia (F); ^***^ = p-value < 0.0001 based on Tukey post-hoc differences. Error bars represent standard error of the mean.

To provide further clarity regarding the variations in viability that we observed in different culture conditions for the malignant cells (MDA-MB-231 and MCF-DCIS.com), we next quantified proliferation by measuring S-phase entry. The results of the proliferation assay paralleled the results of the viability assay for both the MDA-MB-231 and MCF-DCIS.com cells. The percentage of cells entering the S-phase of the cell cycle during the two-hour incubation period with EdU was significantly lower for 400 μM OdDHL-treated MDA-MB-231 cells than for control (0.6% DMSO) cells across all culture conditions (Fig 3A, Fig 3B). The percentage of EdU-tagged cells after the two hour incubation period was also lower for 400 μM OdDHL-treated versus the control cells across all culture conditions for the MCF-DCIS.com cells (Fig 3C, Fig 3D).

**Figure 3.**
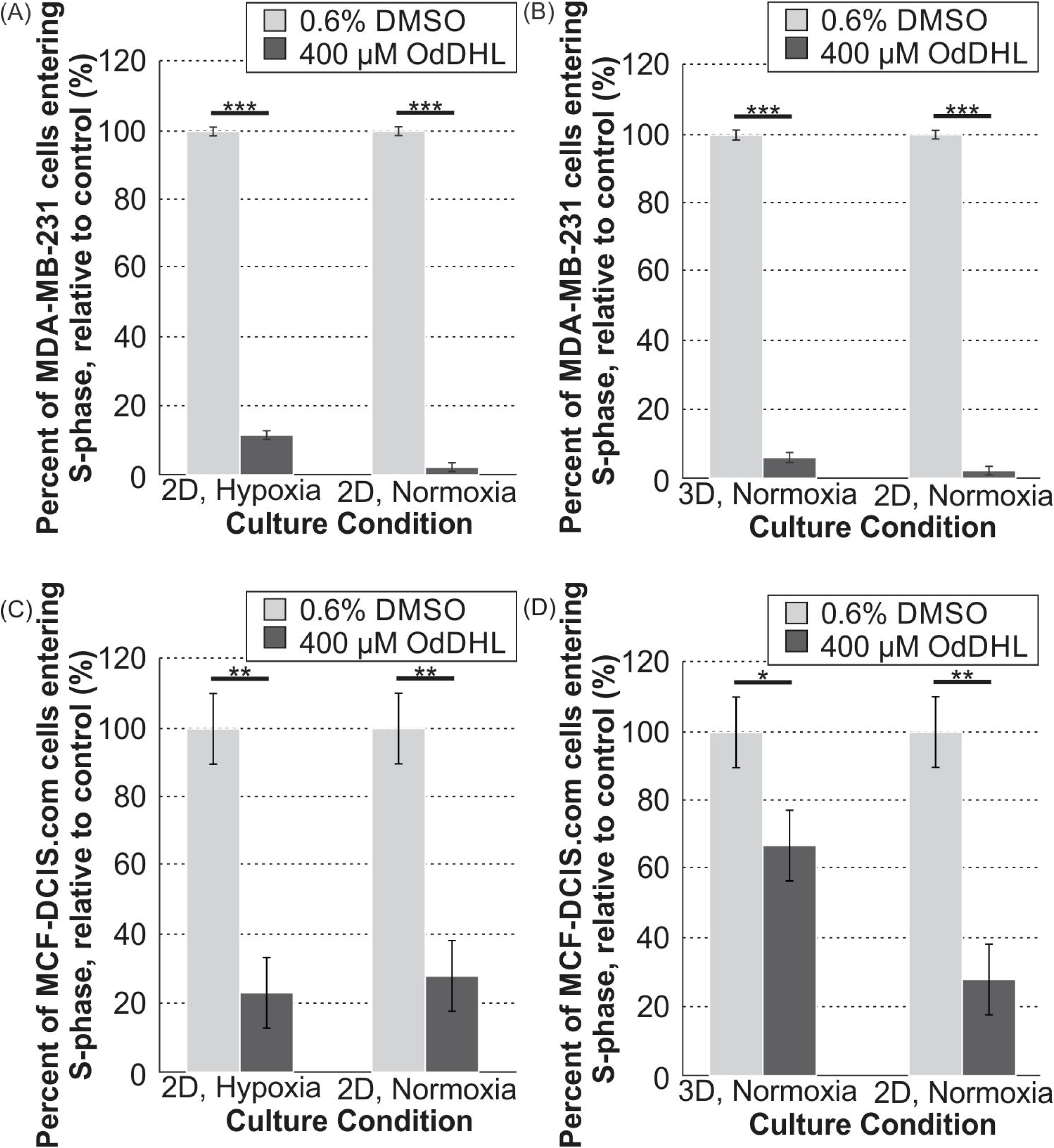
400 μM OdDHL treatment significantly impacts the proliferation rate of MDAMB-231 and MCF-DCIS.com cells in all conditions. Significant differences in the mean percent of MDA-MB-231 cells entering S-phase of proliferation in the 0.6% DMSO and 400 μM OdDHL treatment groups in hypoxia and normoxia in 2D for N = 4 (A) and 3D and 2D in normoxia for N = 3 (B); *** = p-value < 0.0001 based on least mean contrasts. Significant differences in the mean percent of MCF-DCIS.com cells entering S-phase of proliferation in the 0.6% DMSO and 400 μM OdDHL treatment groups in hypoxia and normoxia in 2D for N = 3 (C) and 3D and 2D in normoxia for N = 3 (D); ** = p-value < 0.001, * = p-value < 0.05 based on least mean contrasts. Error bars represent standard error of the mean.

To provide further insights into the viability data, especially in any case where the viability and proliferation trends were not consistent (i.e. the 3D/normoxia condition for MCF-DCIS.com cells), we next quantified apoptosis and necrosis for all cell types. Not surprisingly, the apoptosis/necrosis assay revealed that OdDHL generally induces cell death exclusively for malignant cells. Analysis of the mean gray value (MGV) for the AV-FITC staining showed that apoptosis of MDA-MB-231 cells with 400 μM OdDHL relative to the 0.6% DMSO control in all culture conditions was not significantly increased (Fig 4A). Analysis of the MGV for the PI staining showed significantly increased necrosis of MDA-MB-231 with 400 μM OdDHL treatment, relative to the 0.6% DMSO control, in all culture conditions (Fig 4B). Analysis of MGV for the AV-FITC and PI staining of MCF-DCIS.com cells showed significantly increased apoptosis and necrosis, respectively, only for cells in 2D/normoxia with the 400 μM OdDHL treatment (Fig 4C and Fig 4D). The PI staining of the MCF-DCIS.com cells in the 3D/normoxia showed that these cells had decreased necrosis with 400 μM OdDHL treatment relative to the control, a point to which we will return in the discussion (Fig 4D). The 400 μM OdDHL did not significantly increase apoptosis or necrosis of MCF-10A cells compared to the 0.6% DMSO control (Fig 4E and Fig 4F). Rather, 400 μM OdDHL significantly decreased both MCF-10A apoptosis and necrosis under the 3D/normoxia condition compared to the 0.6% DMSO control (Fig 4E and Fig 4F).

**Figure 4.**
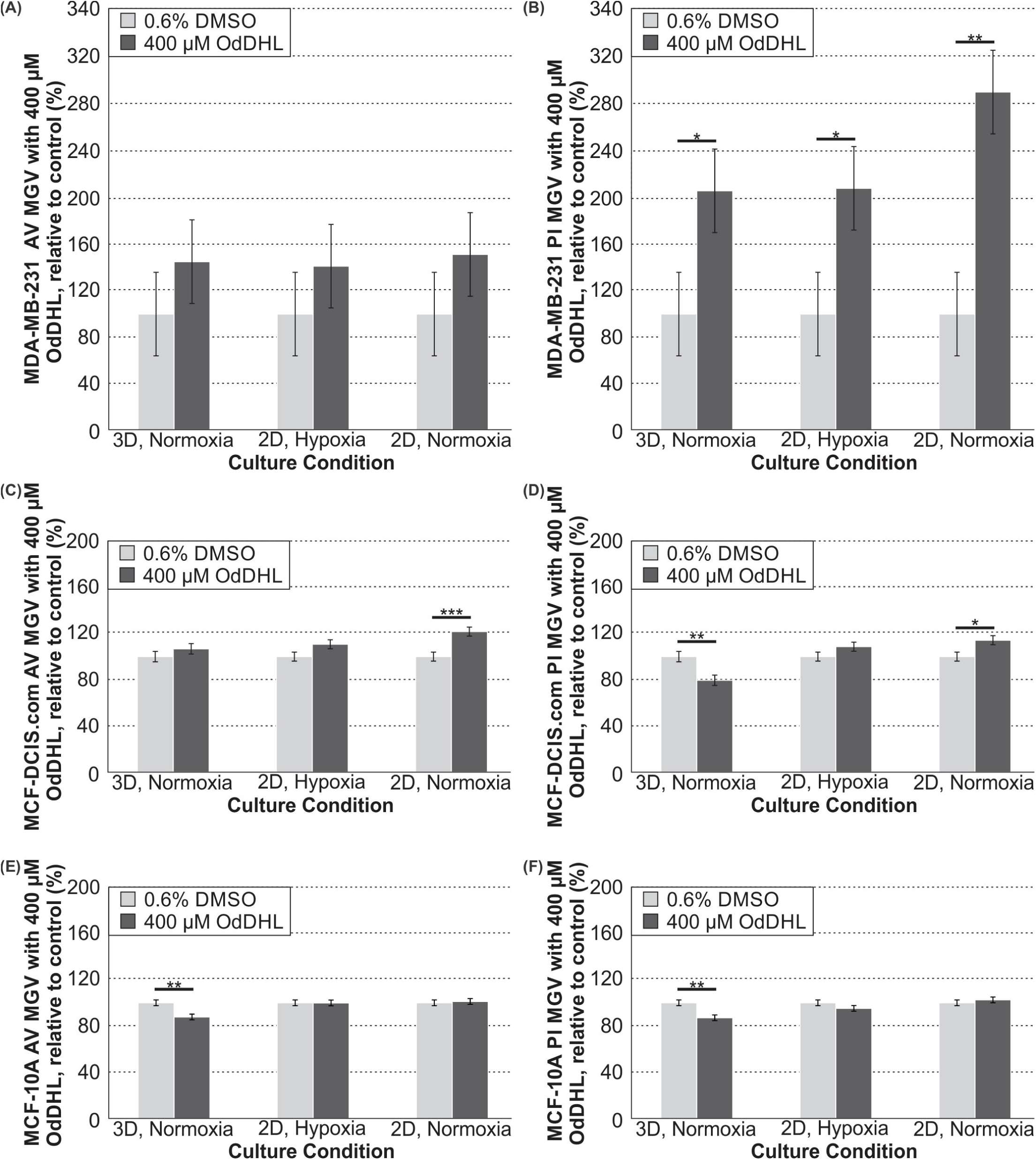
400 μM OdDHL treatment significantly increases necrosis of MDA-MB-231 cells in all conditions. (A) Differences in the annexin V-FITC staining mean gray value (MGV) for MDA-MB-231 cells treated with 400 μM OdDHL, normalized to 0.6% DMSO control for 3D/normoxia, 2D/hypoxia, and 2D/normoxia culture conditions (N = 3). (B) Differences in the propidium iodide staining MGV for MDA-MB-231 cells treated with 400 μM OdDHL, normalized to 0.6% DMSO control for 3D/normoxia, 2D/hypoxia, and 2D/normoxia culture conditions (N = 3); ** = p-value < 0.01, * = p-value < 0.05 based on least mean contrasts. (C) Differences in the annexin V-FITC staining MGV for MCF-DCIS.com cells treated with 400 μM OdDHL, normalized to 0.6% DMSO control for 3D/normoxia (N = 3), 2D/hypoxia (N = 4), and 2D/normoxia culture conditions (N = 4); *** = p-value < 0.001 based on least mean contrasts. (D) Differences in the propidium iodide staining MGV for MCF-DCIS.com cells treated with 400 μM OdDHL, normalized to 0.6% DMSO control for 3D/normoxia, 2D/hypoxia, and 2D/normoxia culture conditions (N = 3); ** = p-value < 0.01, * = p-value < 0.05 based on least mean contrasts. (E) Differences in the annexin V-FITC staining MGV for MCF-10A cells treated with 400 μM OdDHL, normalized to 0.6% DMSO control for 3D/normoxia, 2D/hypoxia, and 2D/normoxia culture conditions (N = 3); ** = p-value < 0.01 based on least mean contrasts. (F) Differences in the propidium iodide staining MGV for MCF-10A cells treated with 400 μM OdDHL, normalized to 0.6% DMSO control for 3D/normoxia, 2D/hypoxia, and 2D/normoxia culture conditions (N = 3); ** = p-value < 0.01 based on least mean contrasts. Error bars represent standard error of the mean.

## Discussion

The analysis of the viability of MDA-MB-231and MCF-10A cells in response to OdDHL confirmed our hypotheses that there would be reduced MDA-MB-231 response to OdDHL treatment in both hypoxia and in 3D while MCF-10A cells would show no such changes. Our 2D/normoxia results are consistent with published literature (21), while our other conditions provide further insight into the interaction of OdDHL with other tumor microenvironment factors. As we anticipated, malignant cells cultured in a 3D environment had the highest relative viability (Fig 1) with OdDHL treatment as compared to cells in 2D. Cells in hypoxia also had increased relative viability with OdDHL treatment as compared to those in normoxia (Fig 1). Thus, we found that OdDHL preferentially affects MDA-MB-231 cells as compared to MCF-10A cells and this effect is blunted by both hypoxia and 3D culture. This result parallels the common finding that tumor cells are more resistant to chemotherapies when studied in 3D and/or hypoxic conditions (27, 31-34), and suggests that these conditions also represent a more physiologically relevant context for the testing of the impact of microbial factors in the tumor microenvironment.

It is interesting to note that, while MDA-MB-231 cells had a decreased viability response when cultured under hypoxia and 3D conditions, the intermediate malignancy MCF-DCIS.com cells did not respond in the same way (Fig 2). MCF-DCIS.com viability was in-between that of the MCF-10A and MDA-MB-231 cells in 2D/normoxia, statistically similar to the MDA-MB-231 response in 2D/hypoxia, and statistically similar to the MCF-10A response in 3D/normoxia (Fig 2). This novel evidence suggests that bacterial factors may affect mammalian cells differently at each stage of cancer progression. It is possible that the response to OdDHL is linked to epithelial-to-mesenchymal transition as it has previously been shown that OdDHL has an effect on cytoskeletal proteins in multiple malignant cell lines (24, 43). OdDHL may be differentially affecting the MDA-MB-231 cells in comparison to the MCF-DCIS.com cells due to its mesenchymal and highly metastatic characteristics; however, much more work needs to be done for confirmation of this hypothesis.

In order to better understand the changes in viability for the breast cancer cells as measured by the alamarBlue™ assay, we next decided to examine the effect of OdDHL on both cell proliferation and apoptosis/necrosis. For MDA-MB-231 cells, it was revealed that OdDHL has two mechanisms to decrease the cell number. We found that OdDHL significantly decreases proliferation of MDA-MB-231 cells in all culture conditions (Fig 3). We then hypothesized that decrease in proliferation alone could not account for the decrease in viability we observed. A second mechanism of action was found in the apoptosis/necrosis assay. We found that necrosis of MDA-MB-231 cells was significantly increased with the 400 μM OdDHL treatment as compared to the 0.6% DMSO control (Fig 4). These two factors worked in concert resulting in the significant viability decrease observed for MDA-MB-231 cells in all conditions.

These effects were not seen in MCF-10A cells and, in fact, both apoptosis and necrosis were reduced in the 3D/normoxia condition while remaining unchanged in all other conditions (Fig 4). The fact that OdDHL triggers necrosis as opposed to apoptosis in malignant cells is not surprising, as bacteria are known to induce necrosis (44). However, the fact that OdDHL may reduce apoptosis and necrosis for non-malignant cells under a stressful microenvironment (i.e. a stiff collagen matrix) is intriguing. More research is needed to understand the selectivity of necrosis induction and, especially, reduction by OdDHL.

Likewise, we found that the primary mechanism responsible for the decrease in MCF-DCIS.com cell number seen with OdDHL treatment was decreased proliferation (Fig 3). While we did not observe decreased viability for MCF-DCIS.com cells in the 3D/normoxia condition, decreased proliferation was observed in all three culture conditions. The apoptosis/necrosis assay shed some light on this seemingly contradictory result. It was found that, as with the MCF-10A cells, there was decreased necrosis for MCF-DCIS.com cells in the 3D/normoxia condition (Fig 4). From these data, we propose that the opposing effects of the decreased proliferation and the decreased necrosis caused by the 400 μM OdDHL treatment countered each other, leading to the invariable response in viability observed. The MCF-DCIS.com cells exhibited increased apoptosis and necrosis in the 2D/normoxia condition, which would have worked in concert with decreased proliferation in this condition leading to the reduced viability observed (Fig 4). Interestingly, we note that the largest reduction in viability for MCF-DCIS.com cells treated with OdDHL was in the 2D/hypoxia condition, although this condition did not result in any significant increases in apoptosis or necrosis. This is different than the observation for the highly malignant MDA-MB-231 cells, whose viability decreased the most under 2D/normoxia. While this may relate to the fact that hypoxia is linked to and predisposes a highly malignant phenotype (45, 46), future mechanistic work will be needed to further clarify these observations. Taken together, these data are indicative that there may be different mechanisms of OdDHL action on cancer viability and these are cell type and microenvironmental condition dependent. While more work with cells representative of varying stages of progression needs to be completed, these preliminary data suggest that OdDHL may be more effective at suppressing the growth of more malignant or advanced breast cancer.

Future studies investigating the possible role of tumor microenvironmental microbiota should establish if, as is suggested here, OdDHL suppresses highly malignant phenotypes. In addition, further experiments should extend the study of non-malignant and pre-malignant cell types to establish if, as seems to be the case for MCF-10A cells, OdDHL increases the survival of these cells in the presence of tumor microenvironmental stressors. Finally, although we show that OdDHL decreases breast viability through inhibition of proliferation as well as cell type and culture condition dependent induction of apoptosis and/or necrosis, future work should establish the cell signaling pathways associated with the differential responses to OdDHL. Li et al. linked the selective action of OdDHL on malignant cell lines to STAT3 activity (21). However, further studies into the mechanism of action of OdDHL with our experimental conditions are required to see if this explanation also extends to our data.

The breast cancer response to OdDHL described above demonstrates that there is significant interaction between bacterial factors and chemical or physical stresses (e.g. hypoxia and stiff ECM) in the tumor microenvironment. Interestingly, in the case of MDA-MB-231 and MCFDCIS.com cells, we demonstrated that malignant cell hypoxia- and adhesion-mediated resistance are not purely associated with chemotherapeutic compounds, but with native microbiome signaling molecules as well. However, it is also notable that, although the MDA-MB-231 cells demonstrated a blunted response to OdDHL in hypoxia and in 3D, there was still a significant response. Thus, future studies into both microbiome-tumor interactions and the therapeutic potential of OdDHL could provide valuable information.

Many open questions remain regarding the physiological or pathological role of the breast resident microbiome. Prior research has established an association between the composition of the breast microbiota and cancer, demonstrating the ability of the abundant bacteria in breast cancer patients to induce double-stranded DNA breaks and/or cell proliferation (9). However, it is not yet known whether these microbiome compositions are a cause or consequence of the malignancies (9). Our data provide an important contribution to this field, demonstrating that even for one quorum-sensing molecule there is a complex breast cell response that is dependent on both cell type as well as microenvironmental context. Many more studies of breast relevant bacteria secreted molecules are needed to establish the impact of the soluble factor crosstalk between the microbiome and breast cancer during the entire course of progression as well as during therapy response. With our data, we have established the importance of performing these investigations in the context of physiologically relevant *in vitro* models. Thus, such investigations would benefit from the incorporation of microenvironmental stressors. More broadly, co-culture experiments investigating soluble factor interactions between breast tissue relevant bacteria and cancer cells in the context of tumor microenvironmental stressors could provide valuable insights into carcinogenesis as well as patient-to-patient variability in disease course and therapy response.

## Conclusions

We showed that the selective effect of OdDHL on the viability of breast cancer cells is significantly mitigated by two key hallmarks of the tumor microenvironment: hypoxia and stiff ECM. It is important to note that, although there was a blunted response to OdDHL treatment for MDA-MB-231 cells in hypoxia and in 3D, there was still a significant decrease in proliferation and viability, and a significant increase in necrosis with treatment. The intermediary response of MCF-DCIS.com may demonstrate a differential role of microenvironmental bacterial factors over the course of breast cancer progression. Thus, further exploration of microbiometumor interactions and OdDHL as a potential therapy is merited.

## Acknowledgements

We would like to thank Sara Peterson for her assistance in experimental preparation. We would also like to thank Dr. Akanksha Kanitkar for her thoughtful discussions, and Dr. Ann Stevens for useful input on the role of OdDHL in bacterial stress response.

